# Bacterial diversity and strain dynamics in the infant respiratory microbiome during the first year of life

**DOI:** 10.64898/2026.05.20.726687

**Authors:** Camila Gazolla Volpiano, Louise M. Judd, Taylor Harshegyi-Hand, Ryan R. Wick, Peter D. Sly, Patrick G. Holt, Merci Kusel, Deborah H. Strickland, Michael Inouye, Kathryn E. Holt, Guillaume Méric

## Abstract

**Background:** Microbial colonisation of the human body begins immediately after birth, with each body site forming a distinct ecological niche. While early gut microbial dynamics and their links to paediatric health have been considerably studied, the bacterial colonisation of the early infant respiratory system remains poorly understood, particularly at the species and strain level.

**Results:** Here, we generated and analysed whole genome sequencing of 925 isolates from six dominant genera in nasopharyngeal samples from 58 healthy infants enrolled in the Childhood Asthma Study (CAS) in Western Australia, collected longitudinally from birth (2, 6, and 12 months old). Our results expanded genomic reference catalogues and uncovered substantial strain-level diversity in dominant infant airway taxa. Plate-sweep metagenomics identified four microbiome profile groups (MPGs) with age-dependent membership, confirming prior observations while enabling high-resolution species and strain analyses. Community maturation was characterised by a shift from early *Staphylococcus aureus* dominance to increased *Moraxella catarrhalis* dominance by 12 months of age, alongside marked temporal changes in prevalence and cohort-level strain diversity.

**Conclusions:** These findings resolve infant nasopharyngeal microbiota composition at the species and strain level, revealing taxon-specific colonisation patterns. By substantially expanding publicly available reference genomes for underrepresented airway taxa, this work also provides a foundation for functional follow-up studies of the early respiratory microbiota and respiratory outcomes.

## Background

The early-life respiratory microbiome assembles rapidly after birth, with initial colonisers detectable within minutes of delivery [1], followed by marked niche differentiation that results in a community typically less diverse than that of the gastrointestinal tract [2,3]. Notably, the nasopharyngeal niche functions as a key ecological reservoir at the host-environment interface, where microbial community structure can modulate mucosal immune tone and influence viral acute respiratory infections [4,5]. The microbial community composition and assembly trajectories of this niche are driven by a complex interplay of perinatal, nutritional, and environmental factors, including delivery mode, breastfeeding, exposure to older siblings or day-care, and antibiotic use [6–10].

Early-life birth cohorts are an important tool to characterise the microbiome development in the respiratory tract and its possible links with future health [9–12]. Leveraging this approach, 16S rRNA gene sequencing has been used to profile the nasopharyngeal microbiome (NPM) in the Childhood Asthma Study (CAS), a Western Australian birth cohort recruited from 1996–1998. NPM communities were characterised during the first year of life in 234 infants from CAS [10] and subsequently over a five-year follow-up [12]. Analysis of this longitudinal data revealed that NPM communities formed discrete microbiome profile groups (MPGs) dominated by *Staphylococcus, Moraxella, Streptococcus, Corynebacterium, Haemophilus*, and *Dolosigranulum*. The maturation trajectory was characterised by early profiles enriched for transient colonisers (particularly *Staphylococcus* and *Corynebacterium*) that, over time, transitioned toward more stable communities dominated by *Dolosigranulum* and *Moraxella* [10]. *Moraxella*, *Haemophilus*, and *Streptococcus* were found to be associated with acute respiratory illnesses (ARIs) and lower respiratory infections (LRIs), as well as increased asthma risk among early-sensitised children. In contrast, *Staphylococcus*, *Dolosigranulum*, and *Corynebacterium* were associated with the absence of respiratory illness [10,12]. Various independent studies have demonstrated similarly reproducible development patterns with consistent associations with respiratory health outcomes. While the colonisation by *Corynebacterium* and *Dolosigranulum* is generally associated with a reduced risk of respiratory illness, *Moraxella*, *Haemophilus*, and *Streptococcus* consistently correlate with an increased risk of infections, recurrent wheeze, and asthma development across diverse populations [9,13–15]. The role of *Staphylococcus* appears more complex and context-dependent, as it is frequently associated with a healthy trajectory and reduced infection risk [9,13] but has also been reported as a risk factor for recurrent wheezing and asthma [11].

The characteristically low microbial biomass and high host DNA content of nasopharyngeal samples have slowed the adoption of shotgun metagenomic approaches in early-life airway microbiome research, which has continued to rely predominantly on 16S rRNA gene sequencing. However, this reliance on genus-level profiling limits functional characterisation, obscures species- and strain-level diversity, and makes it difficult to interpret the inconsistent associations across cohorts. At the same time, inference from metagenomes is constrained by incomplete reference genome collections for several dominant NPM species. There is thus a pressing need for whole-genome sequencing of isolates and shotgun metagenomic approaches at species and strain resolution, both to enable functional experimental follow-up and to inform the design of targeted interventions that may reduce the burden of paediatric respiratory disease.

In this study, we applied a dual approach combining plate-sweep metagenomics with isolate whole-genome sequencing to longitudinal nasopharyngeal samples from infants in the CAS cohort. This allowed us to resolve developmental trajectories of the infant NPM at species and strain resolution, revealing dynamics that differ by taxon. We additionally expanded the public genome catalogue for under-represented airway taxa, and characterised virulence factor and antimicrobial resistance gene repertoires across dominant lineages. More broadly, our study significantly expands the genomic resources and toolkit available to the research community to investigate the putative bacterial effectors of the infant airways and beyond. Our work provides a foundation for identifying the bacterial lineages that shape early-life airway colonisation.

## Results

### Bacterial richness recovered from archival infant nasopharyngeal samples

Our combined sequencing approach characterised the cultivable bacterial composition of archival nasopharyngeal samples collected during periods of health from 58 infants at approximately 2, 6 and 12 months of age (**Table 1**). We generated 142 plate-sweep metagenomes and sequenced 1,036 single colonies. After quality filtering, contamination screening, and representation thresholds (**Methods**), 925 isolate genomes were retained for downstream analyses (**Tables 2, S1**). These 925 genomes comprised 72 distinct species spanning six dominant genera: *Staphylococcus*, *Streptococcus*, *Moraxella*, *Corynebacterium*, *Dolosigranulum*, and *Rothia. Staphylococcus aureus* was the most frequently recovered species (*n*=231 genomes), followed by *Moraxella catarrhalis* (*n*=165), *Corynebacterium pseudodiphtheriticum* (*n*=96), *Dolosigranulum pigrum* (*n*=58), and *Streptococcus pneumoniae* (*n*=46). Five of the six major genera previously reported by 16S rRNA gene surveys of the infant NPM in this cohort [10,12] were recovered here. *Haemophilus* was the notable exception, detected in only 4 isolates and 8 metagenomes despite the inclusion of *Haemophilus*-selective media (HAEM). This likely reflects the known poor viability of *Haemophilus influenzae* under long-term frozen storage relative to other respiratory taxa [16,17]. Given this bias, *Haemophilus* was excluded from subsequent analyses.

**Table 1.** Characteristics of the 58 infants contributing nasopharyngeal samples in this study. Counts are shown as *n* (%). ARI, URI and LRI episodes are cumulative counts of acute, upper and lower respiratory tract infections recorded in the first year of life, respectively. Chronic wheeze was assessed at 12 months of age and refers to persistent wheezing outside of ARI episodes.

| Characteristic | Distribution |
| --- | --- |
| Sex (Male) | 31 (53.4%) |
| Chronic wheeze at 12 months, *n* (%) | 17 (29.3%) |
| ARI episodes (first year), *n* (%) |  |
| 0 | 4 (6.9%) |
| 1–2 | 22 (37.9%) |
| 3–4 | 22 (37.9%) |
| ≥5 | 10 (17.2%) |
| URI episodes (first year), *n* (%) |  |
| 0 | 8 (13.8%) |
| 1–2 | 32 (55.2%) |
| 3–4 | 15 (25.9%) |
| ≥5 | 3 (5.2%) |
| LRI episodes (first year), *n* (%) |  |
| 0 | 21 (36.2%) |
| 1–2 | 33 (56.9%) |
| 3–4 | 3 (5.2%) |
| ≥5 | 1 (1.7%) |

**Table 2.** Taxonomic distribution of 925 bacterial isolate genomes recovered from infant nasopharyngeal samples. For each genus, the total number of isolate genomes and the number of unique infants from whom they were recovered are shown. Species-level assignments under GTDB R214 taxonomy are listed with isolate counts in parentheses and are reported only when more than one isolate is identified. For *S. mitis* and *S. oralis*, GTDB species distinguished by alphabetic suffixes were aggregated into a single species, consistent with recent recommendations [18]. *S. mitis* aggregates *S. mitis*_AQ (*n* = 9), *S. mitis*_AD (*n* = 6), *S. mitis*_I (*n* = 4), *S. mitis* (*n* = 5), *S. mitis*_D (*n* = 3), *S. mitis*_A (*n* = 2), *S. mitis*_AC (*n* = 2), *S. mitis*_AF (*n* = 2), *S. mitis*_BB (*n* = 2), *S. mitis*_BM (*n* = 2), *S. mitis*_K (*n* = 2), and *S. mitis*_P (*n* = 1). *S. oralis* aggregates *S. oralis* (*n* = 5), *S. oralis*_BA (*n* = 3), *S. oralis*_BC, *S. oralis*_I, and *S. oralis*_S.

| Genus | Isolate genomes (*n*) | Infants colonised, *n* (%) | Species identified (*n* isolates) |
| --- | --- | --- | --- |
| *Staphylococcus* | 284 | 81% (47) | *S. aureus* (231), *S. lugdunensis* (14), *S. hominis* (10), *S. capitis* (6), *S. ureilyticus* (6), *S. saprophyticus* (5), *S. pasteurii* (4), *S. epidermidis* (2), *S. pseudintermedius* (2), *S. cohnii*, *S. haemolyticus*, *S. simulans*, and *S. warneri* |
| *Streptococcus* | 207 | 65.5% (38) | *S. pneumoniae* (46), *S. mitis* (40), *S. agalactiae* (23), *Streptococcus* sp. (18), *S. lactarius* (12), *S. oralis* (11), *S. pseudopneumoniae* (7), *S. dysgalactiae* (5), *S. sp001556435* (5), *S. sp903645285* (5), *S. intermedius* (4), *S. sp019448285* (4), *S. parasanguinis_E* (3), *S. salivarius* (3), *S. sp000187445* (3), *S. sp901875575* (3), *S. infantis* (2), *S. parasanguinis_L* (2), *S. pseudopneumoniae_C* (2), *S. anginosus*, *S. pneumoniae_E*, *S. pseudopneumoniae_E*, *S. pseudopneumoniae_G*, *S. shenyangsis*, *S. sp000187745*, *S. sp000314795*, *S. sp902373455*, and *S. sp943912555* |
| *Moraxella* | 184 | 48.3% (28) | *M. catarrhalis* (165), *M. nonliquefaciens* (19) |
| *Corynebacterium* | 159 | 62.1% (36) | *C. pseudodiphtheriticum* (96), *C. accolens* (26), *C. propinquum* (24), *C. sp943914355* (6), *Corynebacterium* sp. (2), *C. coyleae*, *C. kefirresidentii*, *C. minutissimum_A*, *C. otitidis*, and *C. simulans* |
| *Dolosigranulum* | 58 | 37.9% (22) | *D. pigrum* (58) |
| *Rothia* | 33 | 34.5% (20) | *R. sp902373285* (22), *R. mucilaginoso_A* (4), *Rothia* sp. (3), *R. mucilaginoso* (2), *R. dentocariosa*, and *R. terrae* |

### Validation of plate-sweep metagenomics against genome-based, 16S rRNA gene and serological data

Among 902 isolates assigned to species level, MALDI-TOF and GTDB R214 agreed for 803 (89%) after reconciliation of historical NCBI species labels with GTDB taxonomy (**Table S2**). The 99 discordant assignments comprised 18 placeholder GTDB species absent from MALDI-TOF libraries and lacking historical NCBI labels; 35 genus-concordant but species-discordant calls, mainly within the Mitis group of viridans group streptococci (VGS), where MALDI-TOF resolution is limited [18–20], and 46 cross-genus discrepancies, likely reflecting incomplete spectral libraries for under-sampled taxa [21] and possible minority contamination during colony picking (**Methods**). Genome-based assignments were used as the reference for downstream analyses.

Plate-sweep metagenomics recovered a greater community breadth than isolate sequencing, detecting 796 species across 222 genera with a median richness of 16 species per infant visit (range: 2-110), compared with 2 (range: 1-7) by isolate sequencing pooled across media. Nevertheless, per-visit richness estimates were correlated between methods (Spearman’s ρ = 0.55, *P*<0.0001; **Figure S1**).

Compared with matched 16S rRNA gene profiles from the same cohort [10,12], plate-sweep genus-level profiles showed agreement beyond chance for the dominant genus per sample (Cohen’s kappa = 0.32, *P* < 0.0001) and similar ranking of the six dominant genera (median Kendall’s tau = 0.45, **Table S3**). Presence/absence agreement at relative abundance ≥1% was highest for *Moraxella* (Jaccard index=0.55), *Staphylococcus* (0.53) and *Corynebacterium* (0.41), and lower for *Dolosigranulum* (0.31), *Streptococcus* (0.17), and *Haemophilus* (0).

To further validate *S. pneumoniae,* we tested whether carriage during the first year of life was associated with IgG1 responses at 12 months to pneumococcal surface proteins A1, A2 or C, extending prior serological validation in this cohort [10]. Of 57 infants with antibody data, 11 (19%) were seropositive to at least one pneumococcal surface protein (**Table S4**). Detection of *S. pneumoniae* by plate-sweep metagenomics at any visit during the first year was associated with seropositivity (OR=7.9, 95% CI 1.6–46.6, *P*=0.004, Fisher’s exact test), whereas restricting carriage to the 12-month visit alone gave a similar but non-significant effect (OR=5.3, 95% CI 0.7–40.5, *P*=0.054, n=41).

### Microbiome profile dynamics during the first year of life

We classified plate-sweep metagenome taxonomic profiles into four distinct microbiome profile groups (MPGs) based on community composition. Unsupervised clustering resolved four major MPG states dominated by *Staphylococcus*, *Corynebacterium*, or *Moraxella*, alongside a mixed group (**Figure 1A**), mirroring the genera with the highest concordance between NPM profiles generated by plate-sweep metagenomics and 16S rRNA gene profiling. Dominance within these profiles was driven by a small set of prevalent taxa: *S. aureus*, *C. pseudodiphtheriticum* and *M. catarrhalis*. MPG membership varied with infant age (**Table 3**), with *Staphylococcus*-dominated profiles most prominent early in life and *Moraxella*-dominated profiles increasing by 12 months, while *Corynebacterium*-dominated and mixed profiles showed no temporal differences. At the genus level (**Figure 1B**), *Staphylococcus* declined with age, *Moraxella* increased reciprocally, and *Corynebacterium* remained relatively stable. These findings show that plate-sweep metagenomics recapitulates the major age-associated trajectories of the infant NPM previously described in this cohort [10,12], while extending them to species-resolved community structure and enabling downstream strain-level analyses.

**Figure 1.**
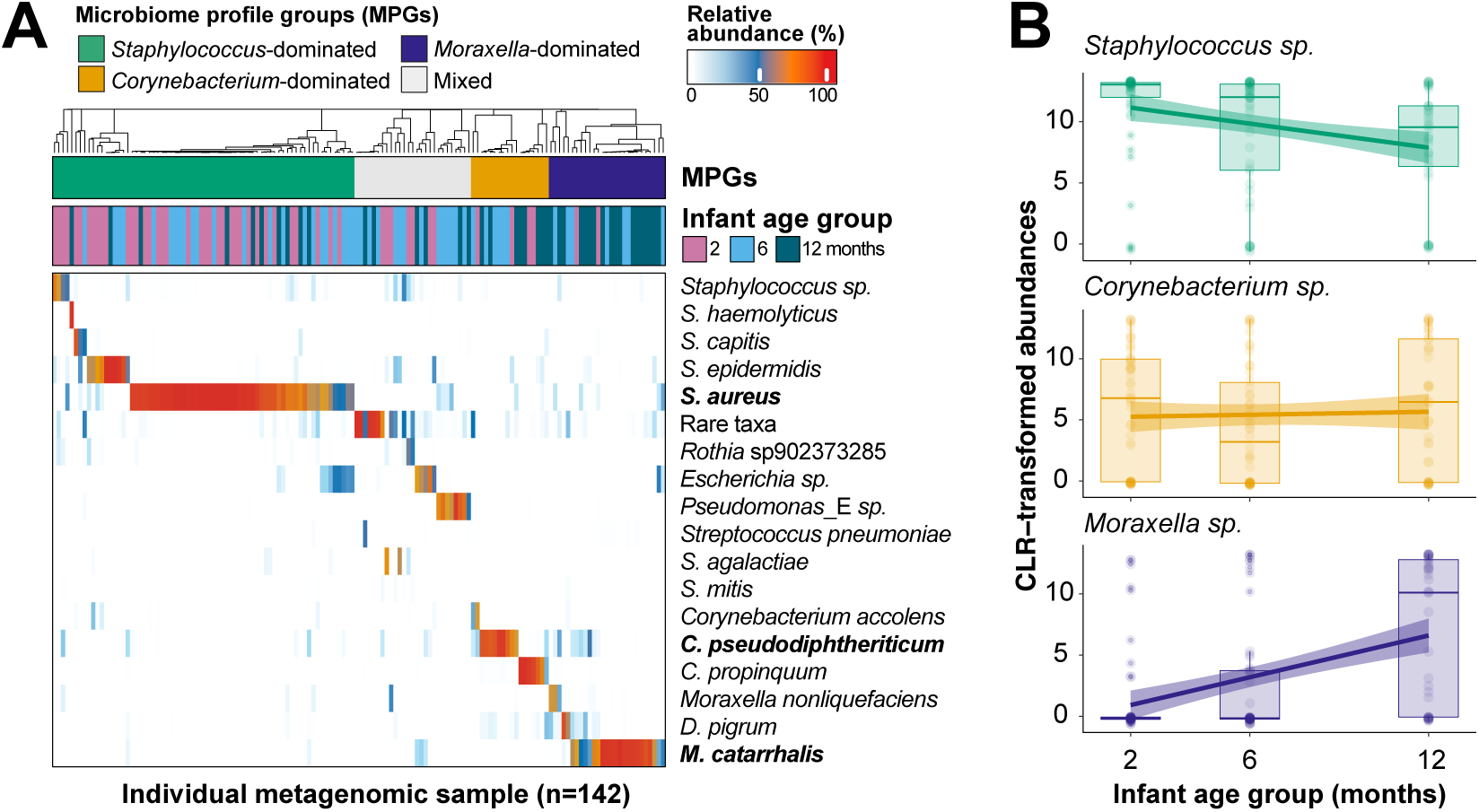
Key bacterial taxa dominate the culturable infant nasopharyngeal microbiota and vary in abundance with age. (A) The heatmap shows the relative abundances of dominant species (mean relative abundance >0.1%, prevalence >20%, and contributing >50% of the relative abundance in at least one sample). Aggregated values are provided for other species from the common genera, as well as for all rare species within each sample. The dendrogram at the top represents complete linkage clustering based on Bray-Curtis distances between samples. Coloured bars indicate the assignment to microbiome profile groups (MPGs) based on this clustering, as well as infant age (2, 6, or 12 months). Dominant species within each MPG, except for mixed, are highlighted in bold. (B) The line plot shows the changes in the centered log-ratio (CLR) abundance of *Staphylococcus* and *Moraxella* genera across the different age groups. The lines represent linear regression fits to the data for each bacterial genus, and the shaded areas represent the 95% confidence intervals around the regression lines.

**Table 3.** Association between infant age and nasopharyngeal microbiome profile groups (MPGs). ORs (95% CIs) for MPG membership at 6 and 12 months relative to 2 months were estimated using logistic GEE clustered by infant. Significant associations (*P* < 0.05) are shown in bold.

| Microbiome profile group (MPG) | 2 months<br>OR (ref.) | 6 months<br>OR (95% CI), $P$ | 12 months<br>OR (95% CI), $P$ |
| --- | --- | --- | --- |
| *Staphylococcus*-dominated | 1 | **0.3 (0.1–0.6), $P = 0.002$** | **0.1 (0.04–0.3), $P < 0.0001$** |
| *Moraxella*-dominated | 1 | 4.5 (0.97–21), $P = 0.054$ | **21.1 (4.3–102), $P = 0.0002$** |
| *Corynebacterium*-dominated | 1 | 1.4 (0.4–4.4), $P = 0.5$ | 2.4 (0.7–7.8), $P = 0.2$ |
| Mixed | 1 | 2.4 (0.9–6.4), $P = 0.1$ | 1.0 (0.3–3.5), $P = 0.98$ |

### Strain-level dynamics of dominant nasopharyngeal taxa

To resolve the strain-level dynamics underlying these species trajectories, we clustered CAS isolate genomes with publicly available references into strain clusters (SCs) using a k-mer-based genome similarity approach [22] at a threshold approximating 99.89% average nucleotide identity (ANI), and used strain-informative k-mers to track these SCs across plate-sweep metagenomes (**Methods**). This showed that the species-level patterns described above were underpinned by distinct dominant strain-level dynamics among dominant NPM taxa (**Figure 2A**).

**Figure 2.**
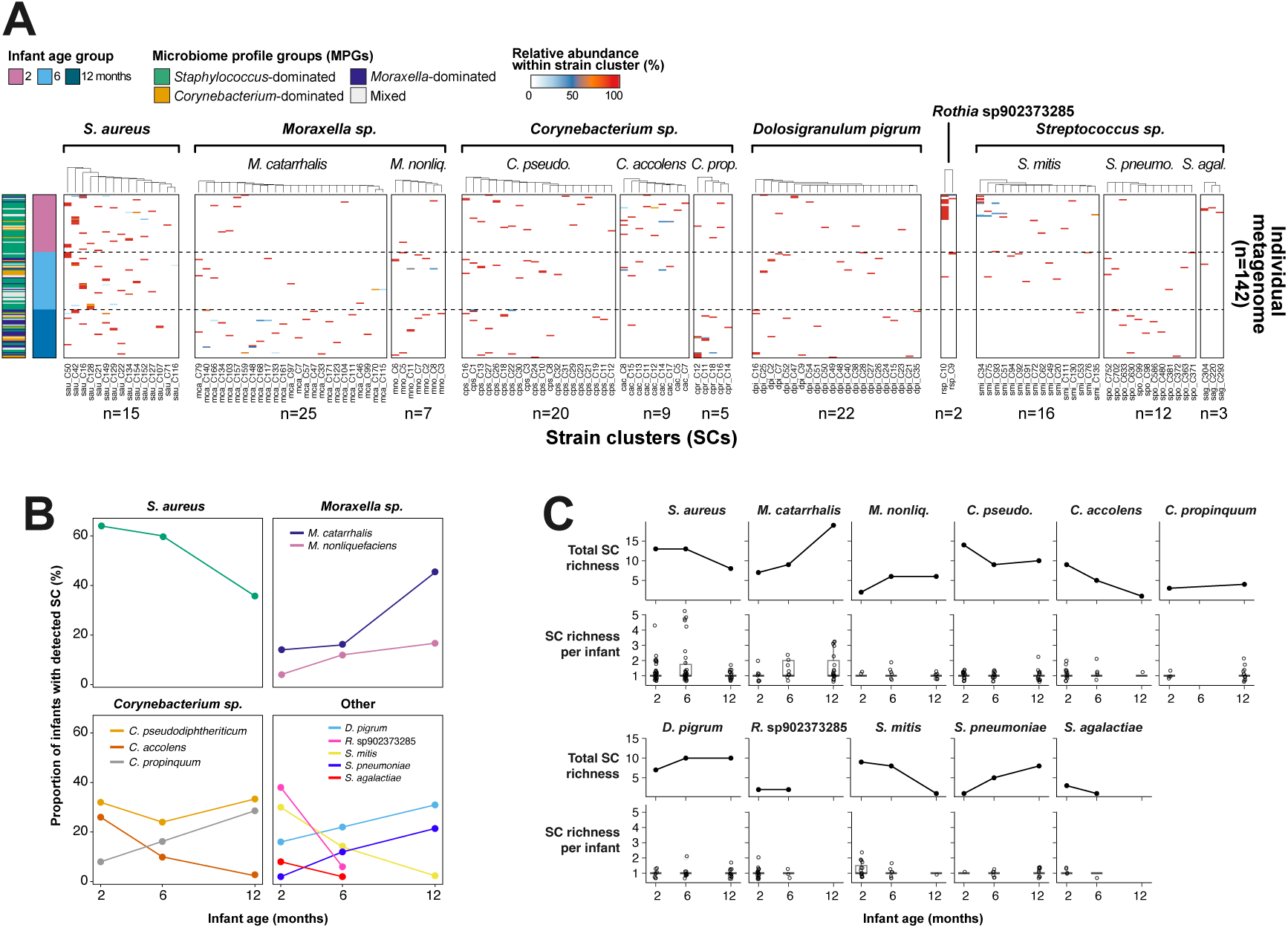
Taxon-specific dynamics of dominant strains in the infant nasopharyngeal microbiome during the first year of life. (A) The heatmap displays the relative abundance of detected dominant strains (∼99.89% ANI) for selected species (panels from left to right: *S. aureus*, *M. catarrhalis*, *C. pseudodiphtheriticum, D. pigrum, S. mitis*, *S. pneumoniae*, *C. accolens, M. nonliquefaciens*, *C. propinquum*, *S. agalactiae*, and *R.* sp902373285). Only strains detected in metagenomes that were defined within a cluster containing a CAS genome are included, as they have stronger evidence of existence. The dendrogram at the top represents complete linkage clustering based on abundance for each species. Coloured bars on the left indicate the assignment to MPGs and infant age (2, 6, or 12 months). Strain IDs are provided at the bottom of the heatmap. (B) Proportion of infants with strains detected via metagenomics, based on the number of infants analysed at different ages: 50 infants at 2 and 6 months, and 42 infants at 12 months (*left*). (C) Variation in strain richness with age, considering all infants (*top*) or each infant individually (*bottom*).

Across taxa, within-host strain richness remained low, with infants typically carrying a single dominant strain (∼99.89% ANI) per species at any given visit; thus, changes in the cohort-wide SC pool mainly reflected shifts in which dominant lineages were circulating across hosts. *S. aureus* was the most prevalent early-life taxon, detected in 64% of infants at 2 months (**Figure 2B**), and its SC pool contracted from 13 dominant lineages at 6 months to 8 at 12 months (**Figure 2C**). *Corynebacterium accolens* and *Streptococcus mitis* showed similar declines in both prevalence and SC pool size over the first year. In contrast, *M. catarrhalis* prevalence rose from 14% at 2 months to 45% at 12 months, while its SC pool expanded from 7 to 19 lineages, indicating increasing prevalence accompanied by increasing strain diversity. *S. pneumoniae* showed the same pattern at lower prevalence. *Moraxella nonliquefaciens* and *D. pigrum* increased in prevalence with a stable SC pool at 6 months, whereas *Corynebacterium propinquum* became more common by 12 months but only showed modest SC pool expansion. By contrast, *Rothia* sp902373285 and *Streptococcus agalactiae* (Group B *Streptococcus*) were detected early, declined by 6 months, and were not detected at 12 months, with both represented by ≤3 SCs across the cohort, consistent with expected vertical transmission of Group B *Streptococcus* at birth [23]. *C. pseudodiphtheriticum* prevalence and SC richness remained comparatively stable across the first year.

These strain-resolved profiles clarify the dynamics of early-life airway NPM colonisation and reinforce prior 16S rRNA gene-based ecological patterns in this cohort, such as early predominance of skin-associated taxa, particularly *Staphylococcus* and *Corynebacterium*, followed by an age-associated rise in *Moraxella*. The early dominance and subsequent turnover of *S. aureus* are consistent with maternal carriage and early-life transmission as important determinants of neonatal airway colonisation [24–26], including reports that many colonised infants carry strains matching those of the mother [27], and that this early seeding event, often occurring around the first month of life, is transient [28]. Conversely, the age-associated increase in both prevalence and strain diversity of *Moraxella* supports predominantly postnatal acquisition of taxa whose usual niche is the respiratory tract, consistent with exposures such as older siblings and day-care attendance [8,29–31].

### Genomic diversity and phylogenetic structure of dominant infant airway species

The addition of CAS isolates to the public genome collection available at the time of this study (April 2024) substantially expanded the genomic landscape of the infant airway microbiota, helping to close diversity gaps for species prevalent in early life but historically under-represented in public repositories (**Table 4**).

**Table 4.**
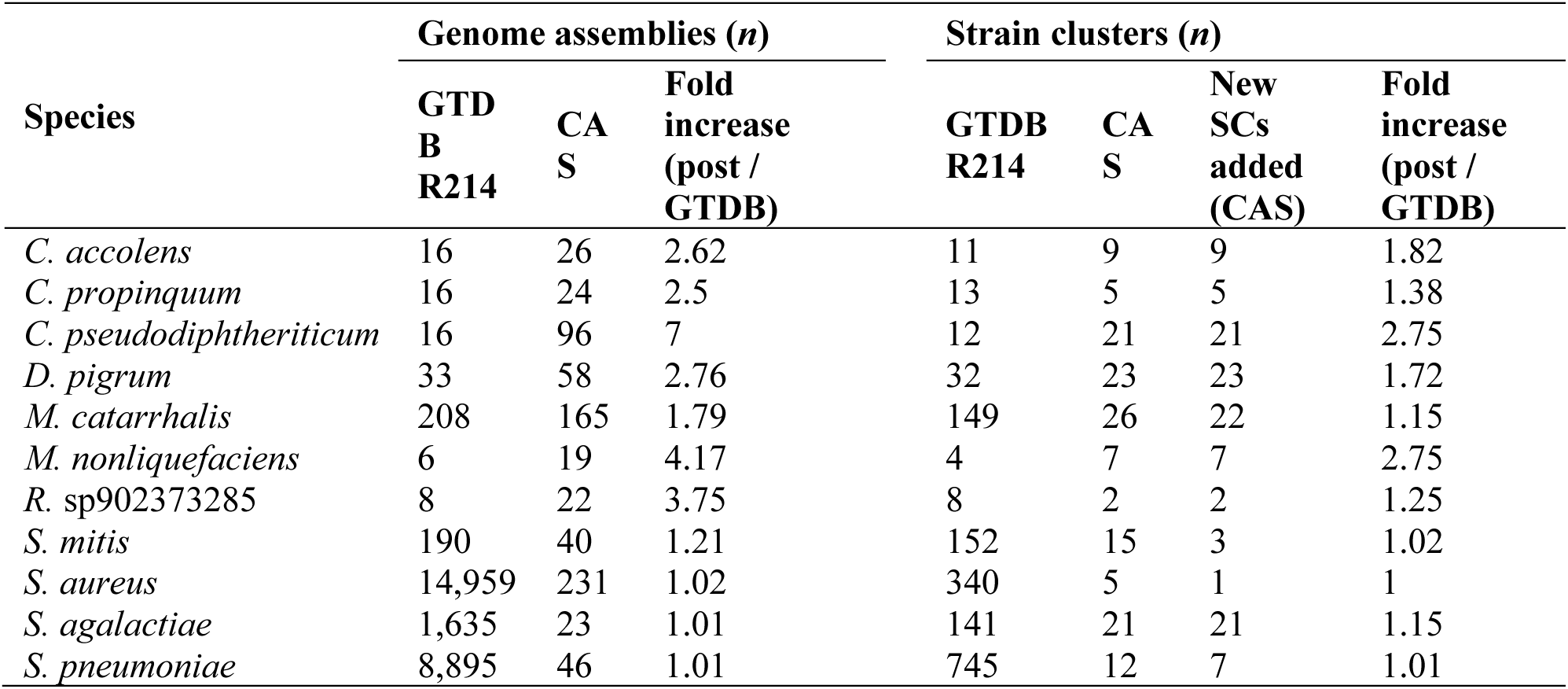
Increase in microbial genomic diversity from infant airways compared to publicly available genomes from GTDB R214 at the time of study. For each species, we report the number of genome assemblies available in GTDB R214 and the number of CAS genomes added, together with the resulting fold increase in total available genomes after adding CAS. We also report the number of SCs in GTDB R214, the number of CAS SCs detected, the number of new SCs added (CAS-only; absent from GTDB), and the corresponding fold increase in SC diversity after incorporating CAS. SC analyses considered all available genomes, except for *S. aureus* and *S. pneumoniae*, for which SC counts were estimated from representative public genome collections (1,248 *S. aureus* genomes and 1,434 *S. pneumoniae* genomes).

The largest gains were observed for *C. pseudodiphtheriticum*, for which available genomic resources increased 7-fold (from 16 to 112 genomes), and species-level strain diversity expanded 2.75-fold. Importantly, this expansion captured novel phylogenetic diversity rather than redundant sampling: all 21 *C. pseudodiphtheriticum* SCs identified in the CAS dataset were absent from public repositories. We also observed substantial novelty for *D. pigrum* and *M. nonliquefaciens*, which contributed 23 and 7 novel SCs, respectively, corresponding to genome assembly increases of 2.76-fold and 4.17-fold. By contrast, the relative contribution of CAS genomes to *S. aureus* and *S. pneumoniae*, both already supported by extensive public collections, was modest in terms of genome counts; however, MLST profiling still identified previously unassigned allelic profiles in the CAS collection (**Table S5**), including six novel *S. aureus* sequence types (*n*=34 genomes) spanning CC30 (*n*=12), CC45 (*n*=11) and CC15 (*n*=1), with a further 10 genomes not assignable to a clonal complex. We additionally identified one novel *S. pneumoniae* ST (2 genomes) and 5 novel *M. catarrhalis* STs (46 genomes).

Across phylogenies of the key taxa dominating the culturable infant NPM (**Figure 3**), CAS isolates either sampled a small fraction of extensive global diversity while spanning multiple lineages, as for *S. aureus* and *S. pneumoniae*, or comprised a large fraction of available genomes, substantially expanding reference representation and revealing multiple SCs. Although *S. pneumoniae* forms recognisable lineages, extensive recombination continuously reshapes core genome sequences and blurs branching structure [32,33]. In our phylogeny, this appeared as a broadly radial pattern, with a similar organisation also evident for *D. pigrum*. A comparable but less well-resolved starburst pattern was observed for *C. propinquum*, *C. accolens*, and *M. nonliquefaciens*, although smaller sample sizes represented these species. *S. mitis* also showed a partially radial core, but with longer branches, suggesting greater deep divergence and clearer substructure. Beyond phylogenomics, capsular serotyping provided a vaccine-relevant axis of *S. pneumoniae* diversity. Because CAS samples were collected before the widespread introduction of pneumococcal conjugate vaccines (PCVs) in Western Australia in 2005, the recovered serotypes provide a baseline snapshot of early-life carriage in this vaccine-naïve infant population. Across 16 infants with detections (**Table S6**), most carriage was driven by classical paediatric serotypes now targeted by PCVs: 81% of detections were serotypes included in the PCV7 (23F, 19F, 14, and 06B) or in the broader PCV13 (19A and 06A). The non-vaccine serotypes 15B/15C and 11A were present as minority constituents in the infant upper airway nearly three decades ago.

**Figure 3.**
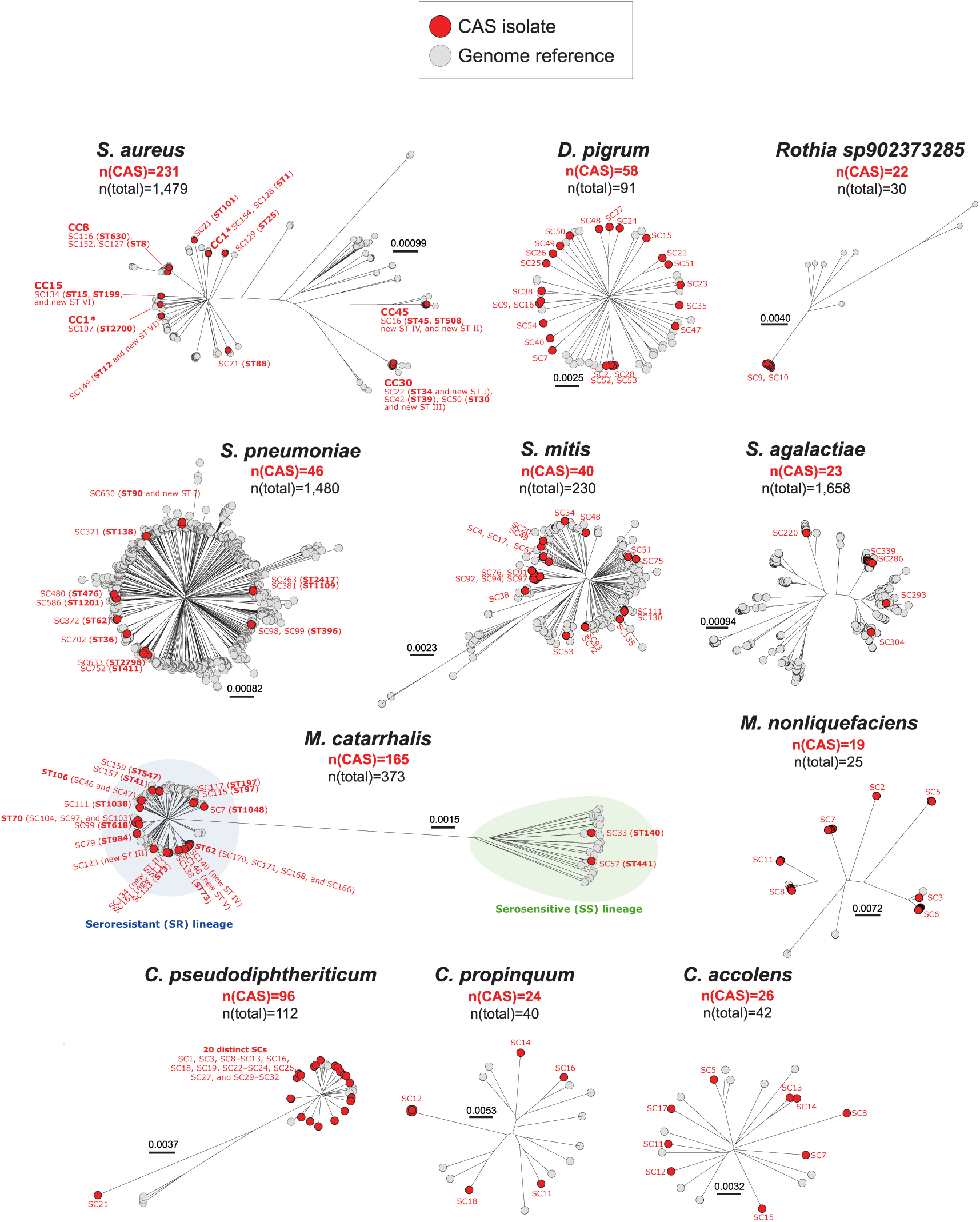
Genetic diversity of isolates from dominant bacterial species from infant nasopharyngeal samples. Neighbour-joining phylogenies were reconstructed from split k-mer-based SNP alignments for *S. aureus*, *D. pigrum*, *R.* sp902373285, *S. pneumoniae*, *S. mitis*, *S. agalactiae*, *M. catarrhalis*, *M. nonliquefaciens*, *C. pseudodiphtheriticum*, *C. propinquum*, and *C. accolens*. Red circles indicate isolates from this study, and grey circles indicate reference genomes from public repositories. The number of genomes included in the trees is written below species names in each panel. CAS isolates on each tree are annotated with their corresponding strain IDs from Figure 2 (including strains not detected in the plate-sweeps but with a genome available), with, when available, the corresponding equivalence in each species MLST schemes from PubMLST.

In contrast to the broadly radial phylogenies of more recombinogenic taxa such as *S. pneumoniae*, *S. aureus* showed a more strongly structured phylogeny. The *S. aureus* STs identified (ST1, ST8, ST12, ST15, ST25, ST30, ST34, ST39, ST45, ST88, ST101, ST199, ST508, ST630, and ST2700) broadly reflect diversity typical of community carriage and were concentrated within a limited set of clonal complexes, including CC1, CC8, CC15, CC30, and CC45. Although some of these STs have been linked to clinically important lineages in certain settings, genome-level assessment of CAS isolates generally did not identify a gene repertoire typical of high-risk clones (see next section).

As expected, *M. catarrhalis* comprised two deeply divergent phylogenetic lineages. Using the recently described *M. catarrhalis* cgMLST scheme and associated LIN code system [34], we found that CAS isolates spanned both the genetically heterogeneous serosensitive (SS) lineage and the conserved/clonal seroresistant (SR) lineage (**Table S7**). Within the SR lineage, the first two levels of the LIN code showed that isolates were distributed across 10 distinct sub-lineages, the largest being 0_39 (*n*=54 out of 165 *M. catarrhalis* genomes), 0_4 (*n*=46), and 0_5 (*n*=22). The SS lineage was rare, restricted to four genomes of ST140 (SC33; LIN 1_31_1; cgST-411) and three genomes of ST441 (SC57; LIN 1_10; cgST-275), each detected at 12 months in a different infant, with one lineage per infant. To assess whether this lineage distribution was consistent across populations, we examined LIN code assignments reported for 1,428 isolates from the Drakenstein Child Health Study [13,34,35], a longitudinal South African birth cohort of 137 children (**Table S8**). The SS lineage was similarly rare in this independent dataset, accounting for 12.5% of *M. catarrhalis* isolates, indicating that SR-lineage dominance is a reproducible feature of early-life *M. catarrhalis* carriage.

Like *M. catarrhalis*, *C. pseudodiphtheriticum* displayed a strongly partitioned phylogenetic structure, although most genomes clustered within a single dominant clade. Notably, SC21 represented the most divergent CAS lineage and was the only strain carrying *cmx* (see next section), a determinant of chloramphenicol resistance in *Corynebacterium* [36]. For *R.* sp902373285, CAS genomes formed a single compact clade on a long, divergent branch. In contrast, CAS *S. agalactiae* genomes were distributed across distinct lineages within a broader diversity background. Specifically, the 23 isolates spanned four STs/CCs, with each lineage tightly linked to a specific capsular serotype (**Table S9**). The collection was co-dominated by ST8 (CC12) expressing serotype Ib (*n*=10) and ST19 (CC19) expressing serotype III-1 (*n*=10). The remaining genomes belonged to CC1, represented by two ST1 isolates expressing serotype V, and the CC17 lineage, represented by a single ST482 isolate expressing serotype III-2, often described as a hypervirulent clone [37].

Collectively, these findings show that our cultivation approach recovered previously unrepresented phylogenetic diversity among dominant infant airway taxa while substantially improving genome representation for historically under-sampled species.

### Virulence and antimicrobial resistance gene profiles across strains and species

We next characterised the CAS isolates by profiling antimicrobial resistance genes (ARGs) and virulence factors (VFs) across genomes within each SC and related these profiles to SC detection across infant ages in plate-sweep metagenomes (**Figure 4, Tables S10, S11**). Across taxa, VF repertoires were generally dominated by a shared core set that was consistently present within SCs, whereas a smaller subset of accessory VFs and ARGs showed strain-specific distributions.

**Figure 4.**
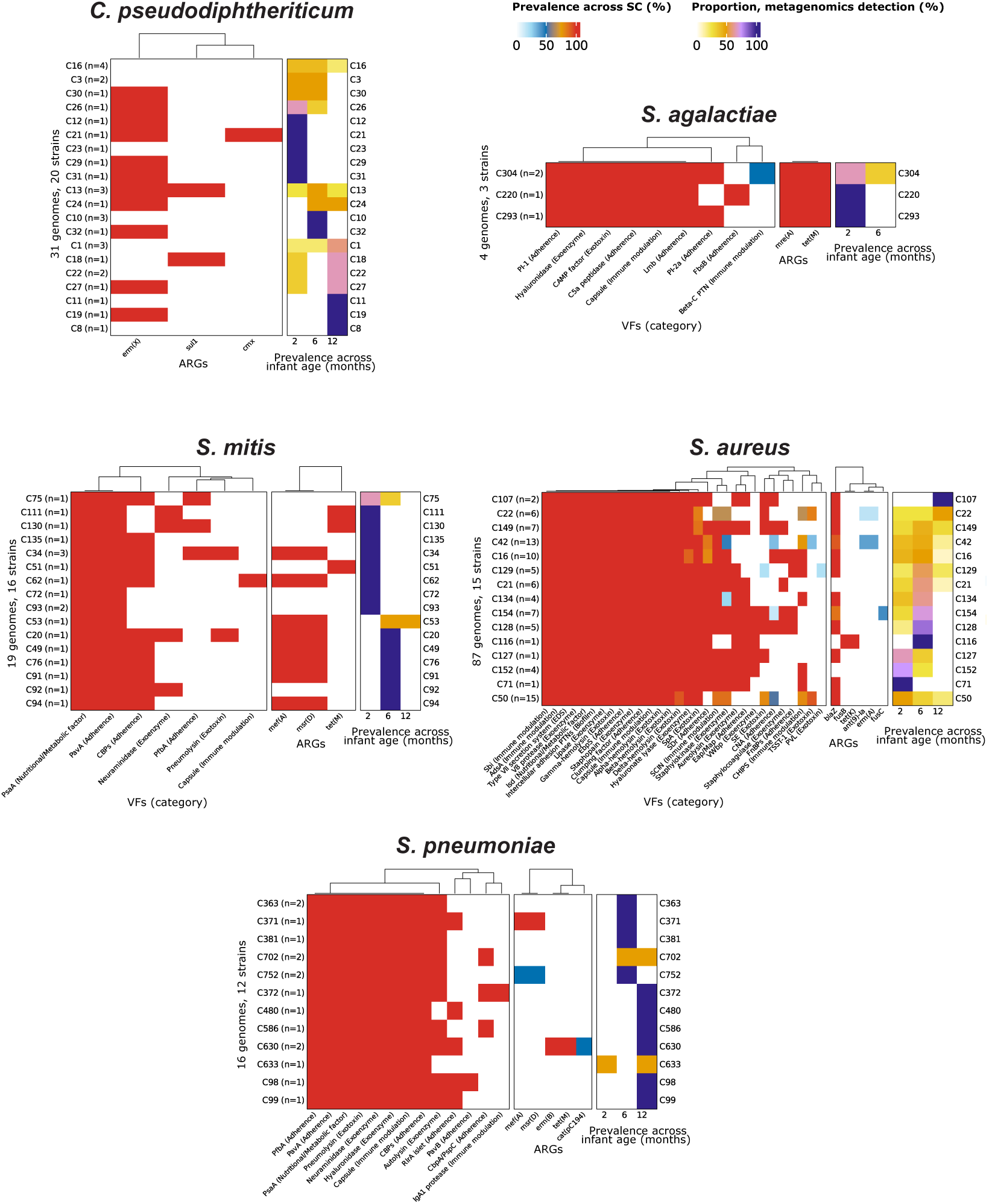
Prevalence of virulence factors (VFs) and antibiotic resistance genes (ARGs) in strains isolated from infant nasopharynx samples. Each row represents a strain cluster detected in at least one cultivation medium and metagenomic sample. The first and second panels display the prevalence of VFs and ARGs across isolates for the identified strains. The third panel indicates the detection of these strains in metagenomic plate-sweeps at different infant ages (2, 6, and 12 months), with colour intensity reflecting the proportion of detections.

Within *S. aureus*, 30 VFs were detected, of which 14 were present in 100% of CAS genomes. Three additional VFs, δ-haemolysin, hyaluronate lyase, and SpA (protein A), were represented across all SCs. Together, these determinants comprised the major haemolysins (α-, β-, γ-, and δ-haemolysins), capsule, multiple immune-evasion factors (AdsA and Sbi), adherence proteins (clumping factor and EbpS), and factors involved in host interaction and persistence, including intercellular adhesion proteins associated with biofilm formation, Isd iron-acquisition functions and several secreted enzymes (staphopain, lipase, and V8 protease). A type VII secretion system was also detected across all CAS *S. aureus* genomes. The widespread presence of these loci likely reflects the conserved *S. aureus* virulence backbone rather than increased pathogenic potential of CAS isolates. Consistent with this, the same factors were also highly prevalent in the 1,248 public genomes used for strain detection (>95% for all listed VFs, except SpA at 85%). In contrast, major toxins showed lineage-specific restriction: Panton-Valentine leukocidin (PVL), a pore-forming toxin associated with severe pleuropulmonary complications in children [38], was identified in only two genomes from SC129 (ST25), both recovered from a single infant at the same 2-month time point. In terms of resistance, we detected no staphylococcal cassette chromosome *mec* (*SCCmec*), indicating no methicillin-resistant *S. aureus* (MRSA) in this community cohort. The β-lactamase gene *blaZ* was the most widespread ARG, present in 85% of CAS genomes and across most SCs, consistent with its high prevalence in public *S. aureus* reference genomes (74.8%).

Within *Streptococcus*, the number of detected core VFs varied by species, with six in *S. agalactiae*, eight in *S. pneumoniae*, and two in *S. mitis*. In *S. agalactiae*, core VFs comprised capsule and hyaluronidase together with adherence-associated factors (Lmb, C5a peptidase, and PI-1) and the CAMP factor. In *S. pneumoniae*, the core VF repertoire also included capsule and hyaluronidase, together with neuraminidase and pneumolysin, the nutrient acquisition factor PsaA, and multiple adherence determinants including PavA, PfbA, and choline-binding proteins (CBPs). By contrast, *S. mitis* had a minimal core VF set limited to PsaA and PavA, both also present in *S. pneumoniae*. ARG profiles also differed across these taxa. All *S. agalactiae* genomes carried *mreA* and *tetM*, whereas ARG carriage in *S. pneumoniae* was restricted to a minority of isolates, with macrolide resistance determinants (*ermB, mefA*, and *msrD*), *tetM,* and *cat(pC194)* confined to a minority of SCs. In *S. mitis*, *mefA* and *msrD* were common (52.6% of genomes), while *tetM* was detected less frequently, in only three SCs.

VF profiling showed no detections for *Moraxella*, *Corynebacterium, D. pigrum,* or *R.* sp902373285. This absence suggests that these isolates either lack canonical virulence markers or carry divergent variants not represented in the database. However, both *Moraxella* species were characterised by the universal presence of *blaBRO-1*. In *Corynebacterium*, ARGs were lineage-specific: *cmx* and *ermX* were identified in a representative *C. accolens* genome (SC12; detected at 2 months), while *cmx, ermX,* and *sul1* were present in *C. propinquum* representatives (SC12 and SC18; detected at 2 and 12 months). In *C. pseudodiphtheriticum*, *cmx, ermX,* and *sul1* were also detected.

Collectively, these results indicate that early-life nasopharyngeal colonisers from CAS largely carry conserved, species-intrinsic VFs and ARGs, while clinically important determinants were rare and lineage-restricted.

## Discussion

We reconstructed the species- and strain-level ecology of the infant nasopharyngeal microbiome during the first year of life using samples collected in the late 1990s. By combining isolate whole-genome sequencing with plate-sweep metagenomics, we recapitulated a major pattern previously described at the genus level by 16S rRNA gene profiling in this cohort, now resolved at a much finer taxonomic scale. Our study also expands the genomic reference landscape for poorly characterised airway bacteria, substantially increasing representation for *C. pseudodiphtheriticum*, *D. pigrum*, *M. nonliquefaciens*, and *S. agalactiae,* and thereby helping offset the bias toward pathogens such as *S. aureus* and *S. pneumoniae* in public repositories of airways isolates.

The shift from *Staphylococcus*-dominated profiles in early infancy to *Moraxella*-enriched communities later in the first year is a defining trajectory of the infant nasopharyngeal microbiome, both reported in this Australian cohort [10,12] and across Europe [9], Africa [13,39,40], Asia [29,41], and the USA [11]. Our analyses suggest that early *Staphylococcus* dominance is driven mainly by widespread early-life colonisation with *S. aureus*, which declines across the first year, likely due to immune maturation and increasing niche competition [7]. The subsequent *Moraxella*-enriched state is driven largely by *M. catarrhalis*, which increased in both abundance and cohort-wide lineage diversity with age, consistent with repeated acquisition of distinct lineages as infant contact networks broaden [10].

In the CAS cohort, *Staphylococcus*-dominated profiles were more common in healthy samples and negatively associated with ARI and LRI, and *Staphylococcus* was also negatively correlated with *Moraxella*, *Haemophilus*, and *Streptococcus*, taxa consistently linked to asthma and wheeze risk in this cohort [10,12]. However, this pattern is not universal: in the COAST cohort from the USA [11], an early *Staphylococcus*-dominant trajectory was associated with increased risk of wheeze, asthma, and early allergic sensitisation. One explanation is that *Staphylococcus*-dominant states are not equivalent across cohorts, but instead differ according to dominant lineages, their virulence and ARG profiles, and the surrounding community and host context. In CAS, our isolate data suggest that *S. aureus* carriage largely reflected low-risk community lineages with limited virulence factor and ARG profiles, consistent with benign colonisation and possibly competitive exclusion of more pathogenic taxa. The sole exception was the detection of an ST25 isolate (SC129) carrying PVL, but this was confined to a single infant at one time point, indicating that such high-consequence virulence determinants were rare in the CAS cohort.

*Moraxella* has been consistently linked to respiratory infections [9,13,42], wheeze [14,43], and asthma [11,15] in cohort studies of the infant airway microbiome. The factors that shift *Moraxella* colonisation from asymptomatic carriage to disease remain poorly defined, but one plausible explanation is differential colonisation by the two deeply divergent *M. catarrhalis* seroresistant (SR) and serosensitive (SS) lineages. SR strains are highly resistant to the human complement and adhere efficiently to respiratory epithelial cells, while SS strains are complement-sensitive and adhere less efficiently [34,44]. Here, the SS lineage was strikingly rare, represented by only two strain clusters, each found in a single infant. In the South African Drakenstein Child Health Study [13,34,35], SS strains were similarly uncommon, comprising 12.5% of isolates. Together, these findings suggest that heterogeneity within the dominant SR lineage, or host and environmental modifiers of *Moraxella* colonisation, are more likely to explain differential respiratory outcomes. The expanded genomic reference collection generated here should help resolve these questions in larger population-based studies.

A key contribution of this study is the expansion of genomic resources for taxa that are common in infancy yet under-represented in public repositories. Respiratory microbiome research has been shaped by sequencing bias toward opportunistic pathogens like *Staphylococcus* and *Streptococcus*, resulting in a scarcity of genomic resources for other dominant airway taxa, including *Moraxella*, *Corynebacterium*, and *Dolosigranulum*. By showing that the CAS isolate set contributes many previously unrepresented strains to GTDB, we highlight this systematic bias and reveal a previously undescribed reservoir of phylogenetic diversity. This reference gap has constrained recent paediatric studies, including efforts to characterise strain-level carriage dynamics in *M. catarrhalis* and *H. influenzae* [45]. In addition, this cohort provides a rare baseline view of the infant nasopharyngeal microbiome before the universal PCV introduction.

Our work also has limitations, including the modest sample size and restriction to Western Australia. We did not directly benchmark plate-sweep metagenomes against uncultured shotgun metagenomes from the same archived aspirates, and the archival nature of the specimens may have imposed biases in culturability and detectability (such as the observed lack of recovered *Haemophilus*). Lower per-strain sequencing depth may also have obscured minor co-colonising strains that drop below the detection threshold. Sampling at only three time points supports developmental inference but cannot resolve rapid strain turnover events. Despite these constraints, the dataset provides a species- and strain-resolved view of infant airway colonisation, and the expanded genome collection for under-represented taxa creates a stronger foundation for future analyses of host-microbe interactions in paediatric respiratory studies.

## Conclusion

This study shows that infant nasopharyngeal microbiome maturation is driven by taxon-specific and strain-specific colonisation dynamics that are not resolved by genus-profiling alone. The transition from early *S. aureus* predominance to later *Moraxella catarrhalis* enrichment was accompanied by contrasting lineage dynamics, while clinically important virulence and resistance determinants remained rare and lineage-restricted. The expanded genome collection generated here provides a species- and strain-resolved baseline for investigating early-life airway colonisation and for interpreting cohort differences in paediatric respiratory outcomes.

## Methods

### Study cohort and sample collection

This study was conducted within the Childhood Asthma Study (CAS), a prospective community-based birth cohort in Perth, Western Australia. The CAS enrolled 244 infants (57% male) at high risk of allergic disease, defined by having at least one parent with a history of doctor-diagnosed atopy (asthma, hay fever, or eczema). Detailed descriptions of the cohort’s design and recruitment have been published previously [10,12,46–48]. In brief, participants were recruited between 1996 and 1998, before the universal introduction of the pneumococcal conjugate vaccine in Western Australia in 2005. Families attended scheduled follow-up visits at approximately 2, 6, and 12 months of age. At each visit, biological samples were collected, and standardised assessments were performed, including anthropometric measurements, clinical examinations, and health questionnaires. Nasopharyngeal samples were obtained during these routine visits only if the child had been free of any acute respiratory illness symptoms for at least 4 weeks, to avoid confounding by recent infections. In addition, participants were visited by a study doctor or study nurse during every episode of parent-reported symptoms of acute respiratory illness (ARI). These episodes were classified as upper respiratory tract infection (URI) or lower respiratory tract infection (LRI) following clinical assessment. The cohort was followed to five years of age, with annual assessment of wheeze and allergy-related symptoms.

### Microbiological cultures

For sequencing of isolate genomes and plate-sweep metagenomes, nasopharyngeal aspirate samples were inoculated onto horse blood agar (BLOO), chocolate agar (CHOC), colistin-oxolinic acid blood agar (COBA), *Haemophilus*-selective agar (HAEM), and *Moraxella*-selective agar (MORA), and incubated at 37°C for up to 7 days at 5% CO_2_. Plates were assessed at 24 hours for growth, and dense growth was collected and diluted/replated to allow for single colony collection. Colonies were picked from each of the five plate types each day for 7 days, focusing on selecting new or novel colonies with different morphologies. MALDI-TOF was used to confirm species identification, and isolates were re-plated for purity and stocking. After 7 days of growth, all plates were swept with Luria broth, and plate-sweep pellets and stocks were collected.

### Genome sequencing, assembly, and taxonomic classification for isolates

A total of 1,036 isolates were subjected to sequencing (**Table S1**). Genomic DNA was extracted using the Illumina DNA Prep Direct Colony Method or the Genfind V3 Genomic DNA Extraction Kit (Beckman Coulter), according to the manufacturer’s protocols. Library preparation was performed using the Nextera DNA Flex Library Prep Kit, using quarter reagents, and the final libraries were sequenced on the Illumina NovaSeq platform, producing 2×150 bp paired-end reads. Reads were assembled using Unicycler v0.5.0 [49] integrated with SPAdes v3.15.5 [50]. Eight isolates failed assembly due to an insufficient number of reads, resulting in 1,028 assembled genome files. Contigs shorter than 500 bp were removed before all downstream quality assessment. Genome completeness and contamination were evaluated using CheckM v1.2.3 [51]. Genomes meeting the following quality criteria were retained for downstream analyses: (i) completeness >50%, (ii) contamination <10%, (iii) quality score (defined as completeness − 5 × contamination) >50, (iv) fewer than 100 contigs, and (v) N50 >5 kb. Applying these criteria, 944 genomes were retained. Taxonomic classification was performed using GTDB-Tk v2.3.0 [52], employing the FastANI v1.32 [53] algorithm and the Genome Taxonomy Database (GTDB) R214 [54]. Species delineation was based on established thresholds: ANI ≥95% and alignment fraction ≥0.5. To ensure adequate representation for comparative analyses, only genomes from genera with more than four isolates (*Corynebacterium, Dolosigranulum, Moraxella, Rothia, Staphylococcus*, and *Streptococcus*) were included. This filtering resulted in a final dataset of 925 genomes derived from 58 infants. Among these infants, genomes were available from 50 infants at 2 months, 43 infants at 6 months, and 40 infants at 12 months of age.

### Retrieval and curation of public bacterial genomes and phylogenetic analysis

Public genome assemblies were retrieved from the NCBI Datasets Genome Package v15.16.2, based on assembly sequence accessions available in GTDB R214 metadata (**Table S9**), for the following taxa: *Corynebacterium accolens* (*n* = 16), *Corynebacterium propinquum* (*n* = 16), *Corynebacterium pseudodiphtheriticum* (*n* = 16), *Dolosigranulum pigrum* (*n* = 33), *Moraxella catarrhalis* (*n* = 208), *Moraxella nonliquefaciens* (*n* = 6), *Rothia* sp902373285 (*n* = 8), *Staphylococcus aureus* (*n* = 14,959), *Streptococcus agalactiae* (*n* = 1,635), *Streptococcus pneumoniae* (*n* = 8,895), and the *Streptococcus mitis* species complex (*n* = 190). For *S. mitis*, GTDB species distinguished by alphabetic suffixes (e.g., *S. mitis*_AC, *S. mitis*_BA, *S. mitis*_BM) were aggregated into a single species, consistent with recent recommendations that ANI-based thresholds inappropriately fragment this species complex and do not necessarily reflect biologically meaningful species boundaries [18]. Genomes listed in GTDB R214 were already filtered using the same completeness and contamination parameters (i, ii, iii) previously applied. We further filtered the *S. aureus* and *S. pneumoniae* public genomes to obtain a representative collection, given resource constraints and the large number of available genomes. Initially, we retained any genome typed with an ST identified among CAS isolates that appeared in <1% of the public collection, yielding 315 *S. aureus* and 56 *S. pneumoniae* genomes. We then refined the selection from the remaining 14,644 *S. aureus* and 8,839 *S. pneumoniae* genomes by applying four criteria: (i) a defined ST, (ii) fewer than 100 contigs, (iii) complete metadata for “ncbi_country,” and (iv) “ncbi_isolation_source” specified as one of the following: blood, nasal, sputum, wound, skin, food, pus, environment, urine, or dairy milk. Finally, we further restricted the resulting dataset from sources with more than 100 genomes by ensuring representation of up to two STs (*S. pneumoniae*) or three STs (*S. aureus*) per source, selected according to quality score. The final dataset included 1,248 public genomes for *S. aureus* and 1,434 for *S. pneumoniae*. For each species, a phylogeny incorporating both CAS and public genomes was reconstructed using Split k-mer Analysis 2 v0.3.7 [55] for alignment (k-mer size = 31) and rapidNJ v2.3.2 for tree construction.

### Isolate typing and capsular characterisation

Whole-genome sequences were typed by multi-locus sequence typing (MLST) using the PubMLST database [56] and the mlst tool v2.23.0 (available at github.com/tseemann/mlst). The following schemes were used for each species: *S. aureus* (saureus: *arcC, aroE, glpF, gmk, pta, tpi, yqiL*), *S. pneumoniae* (spneumoniae: *aroE, gdh, gki, recP, spi, xpt, ddl*), *S. agalactiae* (sagalactiae: *adhP, pheS, atr, glnA, sdhA, glcK, tkt*), and *M. catarrhalis* (mcatarrhalis_achtman_6: *abcZ, adk, efp, fumC, glyBeta, mutY, ppa, trpE*). Available clonal complexes were retrieved from PubMLST and matched to corresponding sequence type profiles for *S. aureus, M. catarrhalis*, and *S. agalactiae*. For *S. aureus*, the staphylococcal cassette chromosome mec (*SCCmec*) types and subtypes, staphylococcal protein A (*spa*) types, and accessory gene regulator (*agr*) groups were determined using staphopia-sccmec v1.0.0 [57], spatyper v0.2.1, and AgrVATE v1.0.2 [58], respectively. *S. pneumoniae* isolates were assigned capsular types using PneumoCaT v1.2.1 [59]. For *M. catarrhalis*, we also used MiST v1.2.0 [60] to assign cgMLST profiles using the recently described scheme and its associated LIN code system [34]. For *S. agalactiae* CAS genomes, capsular serotypes were assigned from genome assemblies using GBS-SBG with the - best parameter [61].

### Profiling of virulence factors and antimicrobial resistance genes

We employed ABRicate v1.0.1 (available at github.com/tseemann/abricate) to identify virulence factors (VFs) and antimicrobial resistance genes (ARGs, **Table S10**) for *C. accolens*, *C. propinquum*, *C. pseudodiphtheriticum*, *D. pigrum*, *M. catarrhalis*, *M. nonliquefaciens*, *R.* sp902373285, *S. aureus*, *S. agalactiae*, *S. mitis*, and *S. pneumoniae*. Searches were performed using the ResFinder database ([62]; 3,077 sequences) and the Virulence Factor Database (VFDB, [63]; 2,507 sequences), with identity and coverage thresholds set to >90% and >60%, respectively. Virulence factors classified as “candidates” were removed from the results.

### Sequencing and taxonomic classification of species in plate-sweep metagenomes

Metagenomic samples were obtained for 142 plate-sweeps derived from 58 infants. Among these infants, metagenomes were available from 50 infants at both 2 and 6 months of age, and from 42 infants at 12 months of age. Metagenomic DNA from plate-sweeps was extracted using the Illumina DNA Prep Direct Colony Method according to the manufacturer’s protocol. Library preparation was performed using the Nextera DNA Flex Library Prep Kit, using quarter reagents, and the final libraries were sequenced on the Illumina NovaSeq platform, producing 2×150 bp paired-end reads. Reads were analysed using Kraken2 v2.1.3 [64] with indices derived from the Human Pangenome Reference Consortium (available at doi.org/10.5281/zenodo.8339732). Unmatched reads were retrieved using utility scripts from KrakenTools (available at github.com/jenniferlu717/KrakenTools). The remaining reads were then assigned to Bacteria and Archaea using a customised 128 GB GTDB R214 index database with the parameter --confidence 0.1. Bracken v2.9 (available at github.com/jenniferlu717/Bracken) was subsequently used to estimate species-level abundances. The minimum number of reads required for classification was set to 250 (-t 250), read length was set to 150 (-r 150), and species was selected as the taxonomic rank to analyse (-l S).

### Antibody-based validation of S. pneumoniae detection in plate-sweep metagenomes

To validate the detection of *S. pneumoniae* by plate-sweep metagenomics, we assessed the association between nasopharyngeal carriage during the first year of life and the development of IgG1 antibodies to pneumococcal surface proteins (A1, A2 or C). Serum samples collected at 12 months of age were assayed for IgG1 antibodies against three pneumococcal surface proteins as previously measured [65]. Infants were classified as seropositive if IgG1 levels to at least one of the three proteins exceeded 2000 ng/mL, and seronegative otherwise. Of the 58 infants in the study, antibody data were available for 57. *S. pneumoniae* carriage was defined as the detection of the species in plate-sweep metagenomes from any of the three sampling visits (2, 6, or 12 months of age). The association between carriage and seropositivity was assessed using Fisher’s exact test. A secondary analysis restricted carriage to detection at the 12-month visit alone (*n*=41 infants with both metagenomic and serological data at this time point).

### Concordance between plate-sweep metagenomes and 16S rRNA gene profiles

To assess whether plate-sweep metagenomic profiles recapitulated the bacterial community previously characterised by a 16S rRNA gene sequencing study [10], we compared genus-level assignments for the six dominant genera (*Corynebacterium*, *Dolosigranulum*/*Alloiococcus*, *Haemophilus*, *Moraxella*, *Staphylococcus*, and *Streptococcus*) across 125 matched infant-visit samples. Three complementary metrics were calculated: (i) the dominant genus per sample was determined in each study and cross-tabulated, with overall agreement and Cohen’s kappa estimated; (ii) within-sample rank concordance was assessed by computing Kendall’s tau across the six genera; and (iii) per-genus presence/absence concordance was quantified using the Jaccard index at a 1% relative abundance threshold.

### Microbiome profile group clustering and association with infant age

Previous CAS microbiome studies have demonstrated clinically meaningful associations with MPGs [10,12]. To establish MPGs, “common species” were defined based on their prevalence and dominance within the dataset, specifically those meeting the following criteria: mean relative abundance >0.1%, prevalence >20%, and contribution of >50% of the relative abundance in at least one sample. All remaining species were then agglomerated into “major genera” (mean relative abundance >0.1% and prevalence >10%) or classified as “rare.” Next, hierarchical clustering using Bray-Curtis dissimilarity as the distance metric was employed to group taxa into MPGs. The optimal number of clusters was determined by maximising the median silhouette value. MPGs were subsequently named according to the dominant genera. For visualisation purposes, *Moraxella* spp. and *Pseudomonas* spp. were removed, and *M. nonliquefaciens*, *S. agalactiae*, *S. pneumoniae*, and *S. mitis* (including any GTDB alphabetic suffix) were added as separate rows; however, for the actual MPG calculations, reads corresponding to these taxa were incorporated into the “rare” category. The MPGs and the relative abundances of taxa used to cluster the samples into MPGs are provided in **Table S12**. We examined how MPG membership changes with infant age by creating dummy variables for each MPG and using 2 months as the reference category. We ran separate logistic regression models using Generalised Estimating Equations (GEE) with an exchangeable correlation structure (R package geepack) to estimate the odds of belonging to each MPG based on age, accounting for within-infant correlation across repeated visits at 2, 6, and 12 months.

### Temporal analysis of strains

StrainScan v1.0.14 [22] was used to identify clusters representing high genomic similarity (“strain clusters”, SCs) among CAS isolates and publicly available genomes (**Table S13**). StrainScan defines SCs by single-linkage hierarchical clustering of genomes using all-vs-all k-mer Jaccard similarity (k-mer size = 31), with a cutoff of 0.95, which corresponds to an average nucleotide identity (ANI) of 99.89%. We also employed StrainScan using its default parameters to detect strain-specific k-mers within each metagenomic sample (**Tables S14** and **S15**). To ensure representative genome selection for each strain, multiple cultivation media were considered. In cases where multiple genomes of the same strain were isolated simultaneously from a single infant, a single representative genome was selected based on the highest quality score, the fewest contigs, and the greatest N50 value.

## Authors’ contributions

Conceptualization: G.M., K.E.H. Cohort establishment and clinical phenotyping: P.D.S., P.G.H., D.H.S., and M.K. Wet lab: L.M.J., T.H.H. Resources (database creation): R.R.W. Formal analysis and visualisation: C.G.V. Supervision: G.M., K.E.H., M.I. Writing of original draft: C.G.V., G.M. Review and editing: All authors. Funding acquisition: G.M., K.E.H., M.I. All authors read and approved the final manuscript.

## Availability of data and materials

The assembled genomes and raw sequencing reads generated in this study have been deposited in the European Nucleotide Archive (ENA) under the project accession number PRJEB111237. Isolate genomes from species for which novel sequence types were identified were submitted to the PubMLST database [56] for curation, where novel alleles and sequence types are assigned official designations (MLST allele combinations from this study are reported in Table S5). All custom code used for analysis is publicly available on GitHub at https://github.com/gazollavolpiano/CAS_2025.

## Declarations

### Ethics approval and consent to participate

The initial CAS protocol for sample collection was approved by the Human Research Ethics Committee of King Edward Memorial and Princess Margaret Hospitals in Western Australia (RE95-17.9 and ECO2-53.9), and written informed consent was obtained from the parents or legal guardians of all participating infants.

### Consent for publication

Not applicable.

